# Neurotoxic Methamphetamine Doses Alter CDCel-1 Levels and Its Interaction with Vesicular Monoamine Transporter-2 in Rat Striatum

**DOI:** 10.1101/2024.07.21.604458

**Authors:** Heli Chauhan, Nick Carruthers, Paul Stemmer, Bernard P. Schneider, Anna Moszczynska

**Author notes:** Correspondence: Anna Moszczynska.

## Abstract

In recent years, methamphetamine METH misuse in the US has been rapidly increasing and there is no FDA-approved pharmacotherapy for METH use disorder (MUD). In addition to being dependent on the drug, people with MUD develop a variety of neurological problems related to the toxicity of this drug. A variety of molecular mechanisms underlying METH neurotoxicity has been identified, including dysfunction of the neuroprotective protein parkin. However, it is not known whether parkin loss of function within striatal dopaminergic (DAergic) terminals translates into a decrease in DA storage capacity. This study examined the relationship between parkin, its substrate cell division cycle related-1 (CDCrel-1), and vesicular monoamine transporter-2 (VMAT2) in METH neurotoxicity in male Sprague Dawley rats. To also assess individual differences in response to METH’s neurotoxic effects, a large group of rats was treated with binge METH or saline and sacrificed 1h or 24h later. This study is the first to show that binge METH alters the levels and subcellular localization of CDCrel-1 and that CDCrel-1 interacts with VMAT2 and increases its levels at the plasma membrane. Furthermore, we found wide individual differences in the responses of measured indices to METH. Proteomic analysis of VMAT-2-associated proteins revealed upregulation of several proteins involved in the exocytosis/endocytosis cycle. The results suggest that at 1h after METH binge, DAergic neurons are engaged in counteracting METH-induced toxic effects, including oxidative stress- and hyperthermia-induced inhibition of synaptic vesicle cycling, with the responses varying between individual rats. Studying CDCrel-1, VMAT2, and other proteins in large groups of outbred rats can help define individual genetic and molecular differences in responses to METH neurotoxicity which, in turn, will aid treating humans suffering from METH use disorder and its neurological consequences.

## Introduction

Methamphetamine (METH) is a highly addictive and powerful central nervous system psychostimulant that induces a feeling of intense euphoria. Therefore, METH misuse is a worldwide health problem. In the United States, there are close to 2,000,000 people who misuse METH, and deaths from METH overdose are rapidly rising ^1,2^. METH overdose is not the only danger to health. METH is highly neurotoxic, and its misuse causes a variety of serious neurological problems and is associated with a risk of developing Parkinson’s disease ^3–5^. Despite numerous clinical trials conducted to date, there is no FDA-approved medication for METH use disorder (MUD) or the neurological problems associated with the disorder. Such medication is particularly needed for people with severe MUD (people who use METH heavily) as they have the most difficulty quitting METH use, are at high risk to overdose on the drug, and suffer the most from neurological consequences of METH neurotoxicity. Our previous studies showed that parkin, a neuroprotective ubiquitin-protein ligase, protected dopaminergic (DAergic) terminals in rat dorsal striatum from METH neurotoxicity ^6^ and might be a potential new drug target in severe MUD ^7^. Others have shown an important role of monoamine transporter-2 (VMAT2) in METH neurotoxicity and MUD ^8–11^. It is not known whether there is an interaction between parkin and VMAT2 in METH neurotoxicity or MUD.

The VMAT2 is a proton pump localized on synaptic monoaminergic storage vesicles and controls intracellular DA levels transporting DA from the cytoplasm into the vesicles in exchange for protons ^12,13^. METH disrupts the proton gradient of the vesicles and stimulates efflux of DA from the vesicles to the cytoplasm ^14,15^. DA released by METH rapidly autoxidizes in the cytoplasm, leading to oxidative stress and terminal damage ^16^. In addition, neurotoxic METH doses rapidly redistribute rat striatal VMAT2 immunoreactivity from synaptic vesicle-enriched, nonmembrane subcellular fractions to a location not retained in the preparation of the synaptosomes, while leaving VMAT2 membrane expression unchanged ^17^ or increased ^18^. This impaired recycling of synaptic VMAT2 vesicles from the plasma membrane to the cytosol suggested a possibility that high-dose METH might render membrane VMAT2 vesicles trapped at the plasma membrane and non-functional while removing other VMAT2 vesicles from the cytosol via retrograde axonal transport. These changes render DAergic terminals with a decreased ability to sequester DA, thus increasing the pro-oxidant effects of the neurotransmitter.

Parkin is an E3 ligase in the ubiquitin-proteasome system that protects DA neurons from diverse cellular insults, including METH and DA-induced oxidative stress ^6,19^. This protection could be partially exerted via the regulation of DAergic neurotransmission. Thus, parkin has been found to regulate DA transporter trafficking between plasma membrane and cytosol *in vitro* ^20^. It is not known whether parkin has a similar effect on VMAT2 under physiological or neurotoxic conditions. The available literature data suggests that parkin may regulate VMAT2 vesicle trafficking via parkin substrates involved in exocytosis-mediated processes, such as trafficking and docking of DA storage vesicles. These substrates are α-synuclein ^21^, α- and β-tubulin ^22^, synaptotagmins IV and 11 ^23,24^, synphilin-1 ^25^, endophilin, dynamin, synaptojanin 1 ^26^, and CDCrel-1 (septin 5) ^27,28^. Without parkin, these substrates would abnormally accumulate. Song and colleagues demonstrated that parkin facilitated the recycling of synaptic vesicles in cultured human DAergic neurons ^29,30^ whereas Feng and colleagues reported that parkin deficit led to an accumulation of synaptic vesicles in neuronal cell bodies in rodent dorsal striatum, due to a parkin deficit-mediated destabilization of microtubules (MTs) ^22^. It is not known whether METH neurotoxicity-induced deficit in striatal parkin levels our group demonstrated ^31^ mediates the redistribution of VMAT2 vesicles away from striatal terminals observed by the aforementioned previous studies ^17,18^.

CDCRel-1 is a filament-forming protein involved in the inhibition of the exocytotic process ^32,33^. Overexpression of CDCrel-1 produces DA neuron-specific cell death ^34^, which in turn, can be prevented by increasing parkin levels ^35^. Both parkin and VMAT2 were shown to be oxidatively damaged and decreased after administration of high METH doses ^36–38^, while the effects of METH on CDCrel-1 have not been studied. Not much is known about the interactions between parkin, VMAT2, and CDCrel-1, under neurotoxic conditions induced by METH. Knowledge of the effects of neurotoxic METH doses on endocytic and exocytic proteins is also limited. We tested a hypothesis that METH-induced parkin deficit increases CDCrel-1 levels in rat striatal neuronal terminals which, in turn, alters the localization of VMAT2 vesicles within the terminals and decreases DA storage.

## Materials and Methods

### Animals

The study employed young adult male Spraque Dawley rats (∼55 days old at the beginning of the study) from Harlan Laboratories (now Envigo, Indianapolis, IN, USA). Upon arrival, animals were pair-housed and maintained under standard environmental conditions in an AAALAC-accredited vivarium. Animals were maintained on a 12h light/dark cycle with *ad libitum* access to food and water. All experiments were approved by the Wayne State University Institutional Animal Care and Use Committee and conducted in compliance with the ARRIVE guidelines.

PO rats were generated by unilaterally overexpressing parkin in the DA nigrostriatal pathway as published previously ^39^. Briefly, rat parkin-encoding AAV2/6 gene transfer vector (AAV2/6-parkin) was microinjected into the left substantia nigra *pars compacta* (1×10^7^ transduction units) whereas non-coding AAV2/6 vector was microinjected into the right one at following coordinates: −5.6 mm (AP) from Bregma, −2 mm (ML) from Bregma, −7.6 mm (V) from the dura according to the Paxinos and Watson’s rat brain atlas. After 3 weeks, the rats were treated with binge METH or saline. The AAV2/6 gene transfer vectors, non-coding AAV2/6 (with a DNA segment cloned upstream of the pgk promoter to adapt the size of vector genome to AAV packaging capacity, AAV2/6) and rat parkin-encoding AAV2/6 (AAV2/6-parkin) were a kind gift from Dr. Bernard Schneider at the Swiss Federal Institute of Technology Lausanne (EPFL), Switzerland.

### Administration of methamphetamine

(+)-Methamphetamine hydrochloride (METH, 8.0mg/kg free base) (Sigma-Aldrich, St. Louis, MO) or saline (1mL/kg) was administered to rats every 2h in four successive intraperitoneal (i.p.) injections. METH neurotoxicity is associated with hyperthermia, which peaks at approximately 1h after each injection. Therefore, core body temperatures of the rats were measured via a rectal probe digital thermometer (Thermalert TH-8; Physitemp Instruments, Clifton, NJ) before the beginning of the treatment (baseline temperatures) and at 1h after each METH or saline injection. Rats were sacrificed by decapitation at 1h or 24h after the last injection of the drug or saline. The brains were removed, METH neurotoxicity-sensitive dorsal striatum (referred to hereafter as the striatum), and the control METH neurotoxicity-resistant cerebellum were dissected out and stored at -80°C until assayed.

### SDS-PAGE and western blotting

Synaptosomal samples (total synaptosomal, membrane/endosomal, and vesicular/cytosolic fractions) were prepared from striatal and cerebellar tissue by differential centrifugation as previously described ^39^. Specifically, tissue pieces were homogenized in 0.5mL 0.32M sucrose with proteases inhibitor cocktail and centrifuged at 800 x *g* for 24min at 4°C. The supernatant was then centrifuged at 22,000 x *g* for 17min at 4°C. The pellet was re-suspended in 150µL ice-cold distilled water and retained as the total synaptosomal fraction. Part of the total synaptosomal fraction was further centrifuged at 22,000 x *g* for 17min at 4°C and the supernatant was retained as vesicular/cytosolic fraction while the pellet was re-suspended in 150µL ice-cold distilled water and retained as the membrane/endosomal fraction. Sample protein concentrations were determined using Bradford protein assay using bovine serum albumin as the standard.

Synaptosomal fractions were subjected to reducing sodium dodecyl sulfate-polyacrylamide gel electrophoresis (SDS–PAGE) under reducing conditions. The amount of 10-20μg proteins was loaded per lane on 4-12% Bis-Tris gels (Life Technologies, Grand Island, NY). Electrophoresis and western blotting were performed as previously described ^40^, utilizing the following primary antibodies: CDCrel-1 (1:1000, overnight at 4°C) (MAB5358; Chemicon International, Temecula, CA); parkin (1:1,000; overnight at 4°C) (Prk8; Cell Signaling Technology, Danvers, MA); actin (1:1,000; 1 h at room temperature) (8H10D10; Signaling Technology, Danvers, MA); alpha-tubulin (1:1000, overnight at 4°C) (sc58668; Santa Cruz Biotech, Santa Cruz, CA, USA), and rabbit polyclonal primary antibody against VMAT2 (1:3000, overnight at 4°C) (NBP1-69750H, Novus Biologicals) as well as appropriate secondary antibodies. Blots were developed using ECL detection and LAS4000 bioimager (GE Healthcare, Piscataway, NJ). Immunoreactivities were quantified using ImageJ software (National Institutes of Health, Bethesda, MD, USA). For standardization across blots, each blot contained all experimental groups. The western blot data was expressed as relative optical density units and normalized to controls on each blot. This approach normalized differences in the development of the blot and across blots.

### Immunohistochemistry

Brain tissue containing substantia nigra *pars compacta* was post-fixed with 4% PFA and processed as previously described ^6^. Citrate buffer antigen retrieval (ThermoFisher, Waltham, MA, USA) was applied to all tissue sections. The sections were incubated overnight at 4°C with the anti-parkin antibody (Prk8; Cell Signaling Technology) and antibody against DAergic marker tyrosine hydroxylase) (AB152, EMD Millipore Corp., Billerica, MA) both diluted 1:100 in the blocking buffer. The sections were then incubated for 2.5 h at room temperature with Alexa Fluor-488 conjugated secondary antibody (Invitrogen, Carlsbad, CA, USA). DRAQ5 (Invitrogen) was used to stain nuclei. The sections were then mounted using Fluoromount mounting medium (Southern Biotech, Birmingham, AL, USA). The immunostaining on each slice (3 sections per slice) was imaged using the Leica TCS SPE-II laser scanning confocal microscope (Leica, Wetzlar, Germany) and averaged per rat.

### Co-immunoprecipitation

Striatal synaptosomal fractions were prepared from rats euthanized at 1h after binge METH. Dynabeads (Life Technologies, Grand Island, NY) were incubated with 2μl of either the rabbit polyclonal VMAT2 primary antibody (NBP1-69750H, Novus Biologicals), mouse monoclonal CDCrel-1 primary antibody (MAB5358; Chemicon International, Temecula, CA), parkin mouse monoclonal primary antibody (Prk8, 1:1000; Cell Signaling Technology), or radioimmunoprecipitation assay (RIPA) buffer (negative control) for 6–12h at 4°C. This was followed by the addition of synaptosomal fractions (100-200µg) and second incubation (12h at 4°C). For all quantitative co-immunoprecipitation studies, the equal protein content of synaptosomal fraction was incubated with beads for each representing group (saline or METH). Following each incubation, the beads were washed three times using phosphate-buffered saline containing 0.02% Tween-20 (PBST). Following immunoprecipitation, Parkin-, CDCrel-1-, and VMAT2-associated proteins were separated from the beads using SDS Tris–Glycine sample buffer (Bio-Rad, Hercules, CA, USA) and heating (10min at 70°C). The supernatants from the beads were run on 4-12% Bis-Tris gels under reducing conditions and subjected to western blot analysis using the CDCrel-1 primary antibody and anti-mouse secondary antibody, as described above.

### Mass Spectrometry

Membrane/endosomal fractions of synaptosomes were incubated with Dynabeads conjugated with anti-VMAT2 antibody and processed as described for co-immunoprecipitation. The resulting eluted samples were run on a gel and the gel was stained with Sypro Ruby stain. Gel pieces were excised from the Sypro Ruby-stained gel, washed, and digested. The gel pieces were first washed with water and 25mM NH_4_HCO_3_/50% acetonitrile for 15min each. The liquid was removed, and the gel pieces were dehydrated in 100% acetonitrile (ACN) and subsequently rehydrated in 50mM NH_4_HCO_3_. After 5min, an equal volume of 100% ACN was added, and the gel pieces were left to incubate in the solution for another 15min. All liquid was removed, the gel pieces were dehydrated in 100% ACN for 5min, the liquid was removed, and the gel pieces were speed vacuum dried for 5min. Next, the following reactions were performed: reduction with 5mM dithiothreitol in 50mM NH_4_HCO_3_, alkylation with 15mM indoleacetic acid in 50mM NH_4_HCO_3_, and overnight digestion with sequencing-grade trypsin (Promega, Madison, WI) in 25mM NH_4_HCO_3_/10% ACN. Following digestion, peptides were extracted from the gel plugs using 50% acetonitrile/0.05% formic acid. The free peptides were then speed vacuumed to dryness and solubilized in 2% ACN/0.1% formic acid.

The peptides were separated by reverse phase chromatography (Acclaim PepMap RSLC C18 column, Thermo Scientific, Waltham, MA), followed by ionization with the Nanospray Flex Ion Source (Thermo Scientific), and introduced into a Q Exactive mass spectrometer (Thermo Scientific). Abundant species were fragmented with high-energy collision-induced dissociation (HCID). Data analysis was performed using Proteome Discoverer 1.4 (Thermo Fisher, Waltham, MA), which incorporated the Mascot algorithm (Matrix Science). The Swiss Prot_2013_03 database was searched against rat protein sequences and a reverse decoy protein database was run simultaneously for false discovery rate (FDR) determination. Secondary analysis was performed using Scaffold 4.2.1 (Proteome Software). Minimum protein identification probability was set at >=95% with 2 unique peptides at >=99% minimum peptide identification probability. Mascot and X! Tandem were searched with a fragment ion mass tolerance of 0.020 Daltons and a parent ion tolerance of 10 parts per million. Carbamidomethylation of cysteine was specified in Mascot and X! Tandem as a fixed modification. Glu->pyro-Glu of the N-terminus, ammonia-loss of the N-terminus, Gln->pyro-Glu of the N-terminus, oxidation of methionine, and acetylation of the N-terminus were specified in X! Tandem as variable modifications. Oxidation of methionine and acetylation of the N-terminus were specified as variable modifications as well. Mass spectrometry processing was done by the Proteomics Core Facility of Wayne State University.

### High-performance Liquid Chromatography

The synaptosomal membrane/endosomal pellet was obtained as described in Chu et al^41^. Briefly, striatal tissue was homogenized in 0.32M sucrose solution. The homogenate was centrifuged at 800xg for 24 min at 4^0^C. The supernatant was centrifuged at 22,000 x *g* for 17min at 4^0^C to obtain the synaptosomal pellet. The pellet was resuspended in 75μl of water and osmolarity was immediately restored by adding an equal volume of pH 7.5 buffer containing 25mM HEPES and 100mM potassium tartrate. The resulting solution was fractionated at 22,000 x *g* for 17min at 4^0^C into membrane/endosomal and vesicular/cytosolic fraction. The membrane/endosomal pellet was re-suspended in perchloric acid (final concentration 0.3N). The resulting solution was then centrifuged at 22,000 x *g* for 30min to obtain the precipitated protein pellet. The supernatant was analyzed for DA content using high-performance liquid chromatography (HPLC) as previously published, using a Shimadzu Prominence HPLC system with electrochemical detection (Shimadzu Scientific Instruments, Columbia, MD, USA).^31^. The protein pellet was resuspended in 1N sodium hydroxide, and its concentration was determined using the Bradford assay. The concentration of DA was quantified by interpolating peak areas relative to those generated by a range of appropriate standards (Sigma Aldrich, St. Louis, MO, USA). The DA values were normalized to protein content in each sample.

### Statistical Analyses

The comparisons made in the study were pre-planned. We established *a priori* the striatum was the METH-affected brain region while the cerebellum was unaffected by METH based on existing knowledge of the effects of binge METH in the rat brain. Samples were assumed to come from populations with the same standard deviations. Two-way or mixed model (if values were missing) repeated measures (RT) ANOVA followed by the Sidak *post hoc* test was performed on body core temperature data. The Greenhouse-Geisser correction was used if there was a lack of sphericity. Differences between control and METH groups were analyzed separately in each synaptosomal fraction by multiple unpaired *t*-tests and the Holm-Sidak method to correct for multiple comparisons. Two-way ANOVA followed by the Holm-Sidak *post hoc* test was employed when data had two variables. Correlations were determined using simple regression analysis and Pearson’s correlation test. Statistical analyses were performed using the GraphPad Prism program (GraphPad Software, San Diego, CA) on the raw data except for the western blotting data. The western blotting data were expressed as relative optical density units on each gel normalized to controls. This approach normalized differences across blots, allowing for standardization across the treatment groups. The data are presented as mean ± standard error (SEM). Statistical significance was set at *p*<0.05.

## Results

### The 4x8mg/kg METH binge elicits variable thermal responses in WT rats

In experimental animals, high doses of METH are known to cause hyperthermia, which is one of the mediators and indicators of METH neurotoxicity ^16,42^. To confirm that METH binge induced hyperthermia in the rats, core body temperature was measured at 1h after each METH or saline injection. Significant increases in core body temperatures were observed in all METH-treated rats as compared to saline controls (two-way RT ANOVA followed by the Sidak *post hoc* test) (Fig.1a, b). All WT rats reached 39°C at least once. There was a significant main effect of the treatment (METH *vs.* saline) in both experimental cohorts i.e. rats sacrificed at 1h and those sacrificed at 24h after the last METH injection (1h cohort: treatment *F*(1,28) = 52.6, *p*<0.0001, time (*F*(2.42,67.8) = 10.7, *p*<0.0001, and treatment *x* time interaction (*F*(4,112) = 16.2, *p*<0.0001; 24h cohort: treatment *F*(1,16) = 60.6, *p*<0.0001), time (*F*(2.42,37.6) = 5.99, *p*<0.01, and treatment x time interaction (*F*(4,62) = 11.6, *p*<0.0001)). It was observed that despite the same METH regiment the thermal responses to METH were highly variable. The METH-treated rats could be divided into two groups depending on the severity of hyperthermia: those with high hyperthermia (HH) and those with low hyperthermia (LH) (Fig.1c, d). Significant differences in core body temperatures were observed between saline and METH treatments as well as between the subgroups (1h cohort: treatment *F*(2,27) = 106, *p*<0.0001, time (*F*(3.15,83.5) = 53.1, *p*<0.0001, and treatment *x* time interaction (*F*(8,106) = 29.3, *p*<0.0001; 24h cohort: treatment *F*(2,15) = 61.1, *p*<0.0001), time (*F*(3.31,49.7) = 23.5, *p*<0.0001, and treatment x time interaction (*F*(8,60) = 18.3, *p* < 0.0001)). Areas under the curve (AUC) were calculated for all METH-treated rats.

**Figure 1.**
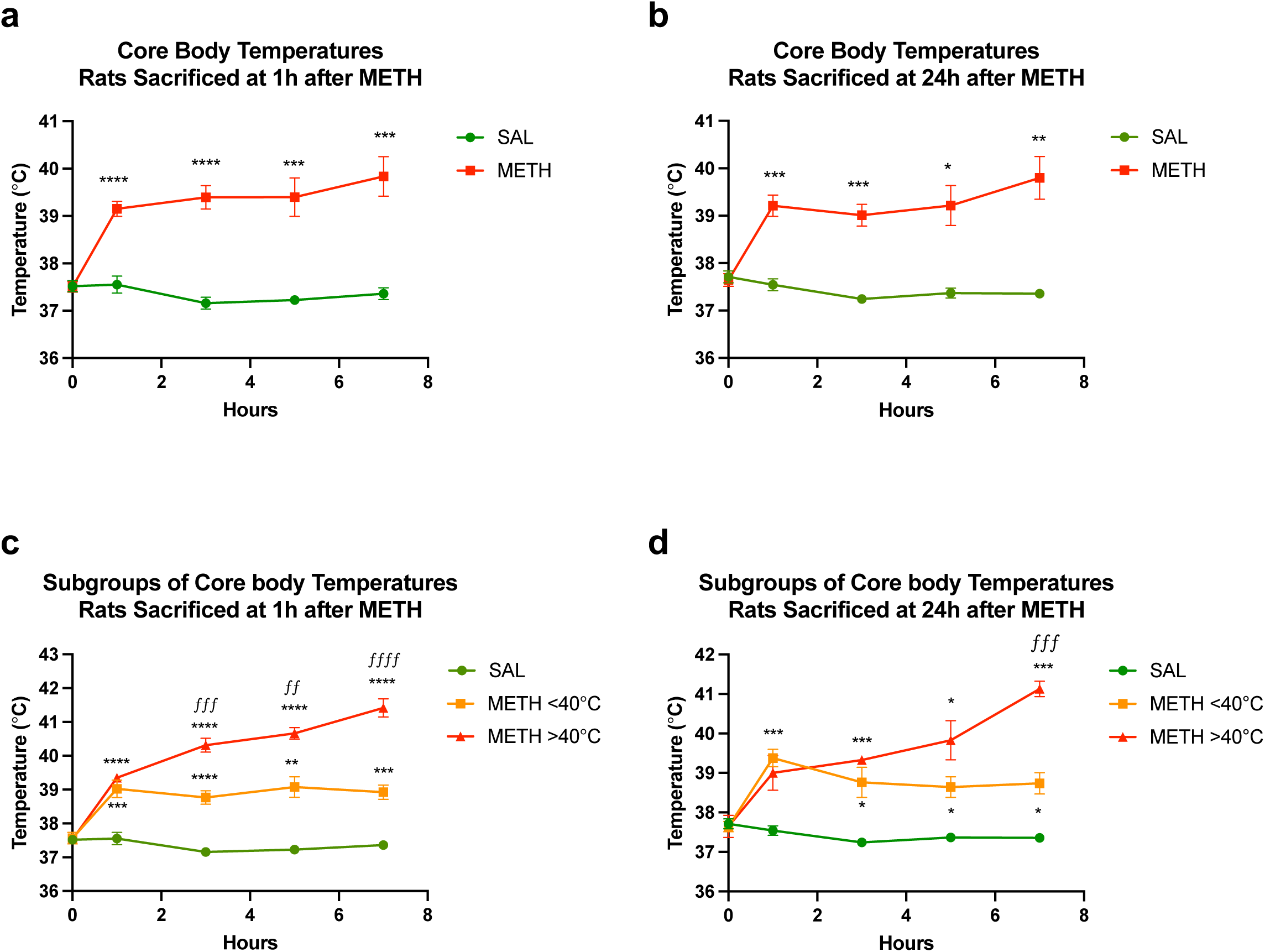
The 4x8mg/kg METH binge elicits variable thermal responses in WT rats. Core body temperatures (°C) were measured before the first METH or saline injection and 1h after each METH or saline injection. Core body temperatures of rats euthanized at 1h **(a)** or 24h **(b)** after METH or saline. **(c, d)** The METH-treated rats were divided into two groups depending on the severity of hyperthermia: those with high hyperthermia and those with low hyperthermia (average of 4 temperature readings >40°C and <40°C, respectively) and euthanized at 1h **(c)** or 24h **(d)** after METH or saline. Significant differences between saline and METH rats: \**p*<0.05, \*\**p*<0.01, \*\*\**p*<0.001, and \*\*\*\**p*<0.0001. Significant differences between HH and LH rats: *^ƒƒ^p<*0.01, *^ƒƒƒ^p<*0.001*, ^ƒƒƒƒ^p<0.0001.* Values are expressed as mean ± SEM. Abbreviations: METH, methamphetamine; SAL, saline.

### The effects of 4x8mg/kg METH binge on α-tubulin and β-actin immunoreactivity in striatal synaptosomal fractions

There is surprisingly little literature on the effect of METH on the cytoskeleton. The available data suggests that METH can alter both MTs and actin filaments ^43^. Therefore, we compared β-actin and α-tubulin immunoreactivities in all striatal synaptosomal fractions at 1h and 24h after the last dose of METH *vs.* saline. β-Actin immunoreactivity was significantly decreased in all three synaptosomal fractions at 1h after METH (total fraction: -17%, membrane/endosomal fraction: -19%, and vesicular/cytosolic fraction: -26%, *p*<0.05, multiple unpaired *t*-tests with Holm-Sidak correction for multiple comparisons, n=7-11) (Fig.2a). No significant changes were detected in β-actin immunoreactivity at 24h after METH (Fig.2b). No statistically significant changes were detected at 1h or 24h after METH in the levels of α-tubulin; however, individual variability in its immunoreactivity was observed in total and membrane/endosomal synaptosomal fraction from binge METH-treated rats at 1h time point (Fig.2c, d), thus suggesting that METH influences α-tubulin levels differently in different rats. Consequently, to control for METH effects on the cytoskeleton, we used total protein levels (measured by Ponceau S dye) as a loading control in subsequent analyses of immunoreactivities on western blots.

**Figure 2.**
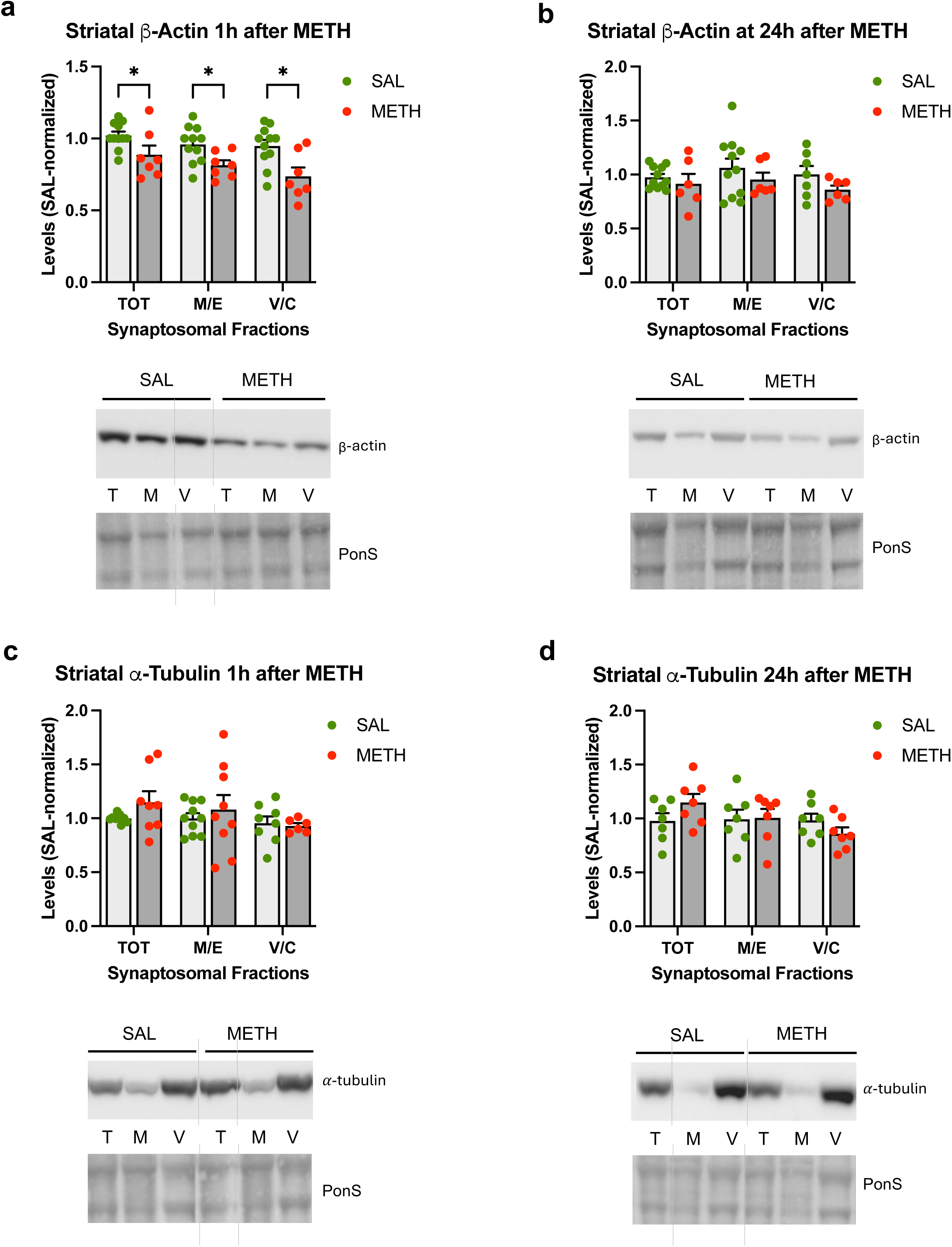
The effects of 4x8mg/kg METH binge on α-tubulin and β-actin immunoreactivity in striatal synaptosomal fractions. Immunoreactivity of β-actin **(a, b)** or α-tubulin **(c, d)** in total (TOT or T), membrane/endosomal (M/E or M), and vesicular/cytosolic (V/C or C) synaptosomal fraction of the striatum in rats that were euthanized at 1h **(a, c)** or 24h **(b, d)** after the last dose of saline or METH. Immunoreactivity of in total (T), membrane/endosomal (M), and vesicular/cytosolic (V) synaptosomal fraction of the striatum in rats euthanized **(a)** at 1h or **(b)** 24h, after the last dose of saline or METH. \**p*<0.05, n=7-11. Values are expressed as mean ± SEM. Vertical grey lines show where the blot was cut for rearrangement. Abbreviations: METH, methamphetamine; SAL, saline; PonS, Ponceau S.

### The effects of 4x8mg/kg METH binge on parkin immunoreactivity in striatal synaptosomal fractions

We previously determined that 4x10mg/kg METH decreased parkin immunoreactivity in total striatal synaptosomal fraction at 1h and 24h after the last dose of the drug, with parkin immunoreactivity returning to the baseline by 48h ^38^. In this study, 4x8mg/kg METH did not significantly decrease parkin immunoreactivity in synaptosomal fractions compared to saline controls; however, a trend toward statistical significance was observed in the total and vesicular/cytosolic synaptosomal fraction (-18% and -20%, respectively, *p*=0.09, unpaired multiple *t*-tests with Holm-Sidak correction for multiple comparisons, n=10-17) at 1h after the last METH dose (Fig.3a). No significant changes in parkin levels (or trends toward statistical significance) were detected at 24h after METH compared to saline controls (Fig.3b). As with core body temperature, high variability in parkin immunoreactivity in METH-treated rats was observed at both time points (Fig.3a, b). When the HH and LH rats from the 1h group were assessed separately, a significant deficit in parkin immunoreactivity was detected in the LH rats in all synaptosomal fractions (two-way ANOVA followed by Holm-Sidak *post hoc* test; main effect of treatment: *F*(1,52)=15.74, *p*<0.001; total fraction: -23%, *p*<0.05; membrane/endosomal fraction: -28%, *p*<0.01; vesicular/cytosolic fraction: -23%, *p*<0.05), with HH rats displaying unchanged parkin immunoreactivity (Fig.3d). However, there was no significant correlation between core body temperature and parkin immunoreactivity in synaptosomal fractions, indicating an influence of additional factor(s) on parkin immunoreactivity in the METH-treated rats. In METH neurotoxicity-resistant cerebellum, a significant increase in synaptosomal parkin was detected compared to saline controls (Suppl. Fig.S1).

**Figure 3.**
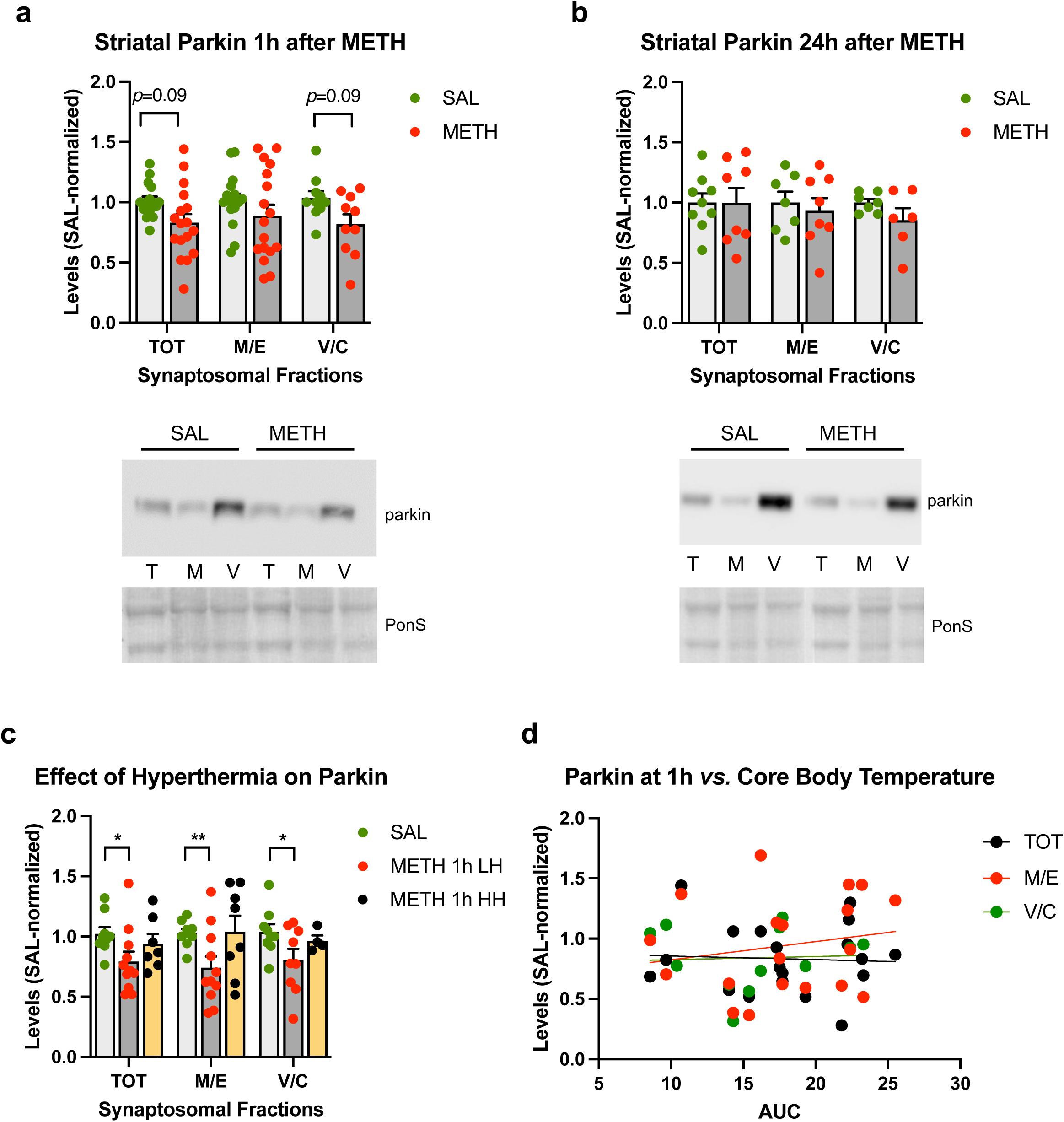
The effects of 4x8mg/kg METH binge on parkin immunoreactivity in striatal synaptosomal fractions. Immunoreactivity of parkin in total (TOT or T), membrane/endosomal (M/E or M), and vesicular/cytosolic (V/C or C) synaptosomal fraction of the striatum in rats euthanized at 1h **(a)** or 24h **(b). (c)** Immunoreactivity of parkin in striatal synaptosomal fractions in high hyperthermia (HH) and low hyperthermia (LH) rats. **(d)** Correlations of parkin immunoreactivity in synaptosomal fractions with core body temperature (area under the curve AUC) of rats sacrificed 1h after the last METH dose. \**p*<0.05, \*\**p*<0.01, n=10-17. Values are expressed as mean ± SEM. Abbreviations: METH, methamphetamine; SAL, saline; PonS, Ponceau S.

### The effects of 4x8mg/kg METH binge on VMAT2 immunoreactivity in striatal synaptosomal fractions

Previous studies have demonstrated that 4x10mg/kg METH rapidly redistributed rat striatal VMAT2 immunoreactivity from the vesicular/cytosolic synaptosomal fraction to a location not retained in the preparation of the synaptosomes while leaving VMAT2 membrane expression unchanged ^17^ or slightly increased ^18^. Specifically, decreases in partially glycosylated VMAT2 (∼55 kDa) were reported in the vesicular/cytosolic fraction at 1h after METH, with membrane VMAT2 levels reported to be increased by one study and not changed by the other ^17,18^. A decrease in glycosylated VMAT2 (∼68 kDa) in vesicular/cytosolic fraction at 1h and decreases across all synaptosomal fractions at 24h after METH has been reported by the Yamamoto group ^44,45^. Of the 3 isoforms of VMAT2, our anti-VMAT2 antibody easily identified the partially glycosylated form (∼55 kDa) and the glycosylated form (68-75 kDa) of the protein; the non-glycosylated VMAT2 (45 kDa) was detected at low levels (Fig.4). At 1h after METH, we did not detect significant changes in VMAT2 immunoreactivity in synaptosomal fractions as compared to saline controls (Fig.4a). At 24h after METH, a significant decrease in VMAT2 immunoreactivity was detected in the vesicular/cytosolic fraction (-34%, *p*<0.05, unpaired multiple *t*-tests with Holm-Sidak correction for multiple comparisons, n=9-11) in METH-treated rats relative to saline-treated rats (Fig.4b). As with individual parkin levels, high variability in individual synaptosomal VMAT2 immunoreactivities was observed after METH binge at both time points, 1h and 24h. Subgrouping VMAT2 values from the 1h METH group according to the levels of hyperthermia did not produce significant changes in VMAT2 immunoreactivity as compared to the corresponding saline group (Fig.4c). There was no significant correlation between core body temperature and VMAT2 immunoreactivity in any of synaptosomal fractions in METH-treated rats (Fig.4d); however, the correlation between core body temperature and membrane/endosomal VMAT2 displayed a strong trend toward statistical significance (*p*=0.055, R^2^=0.21, Pearson’s correlation test).

**Figure 4.**
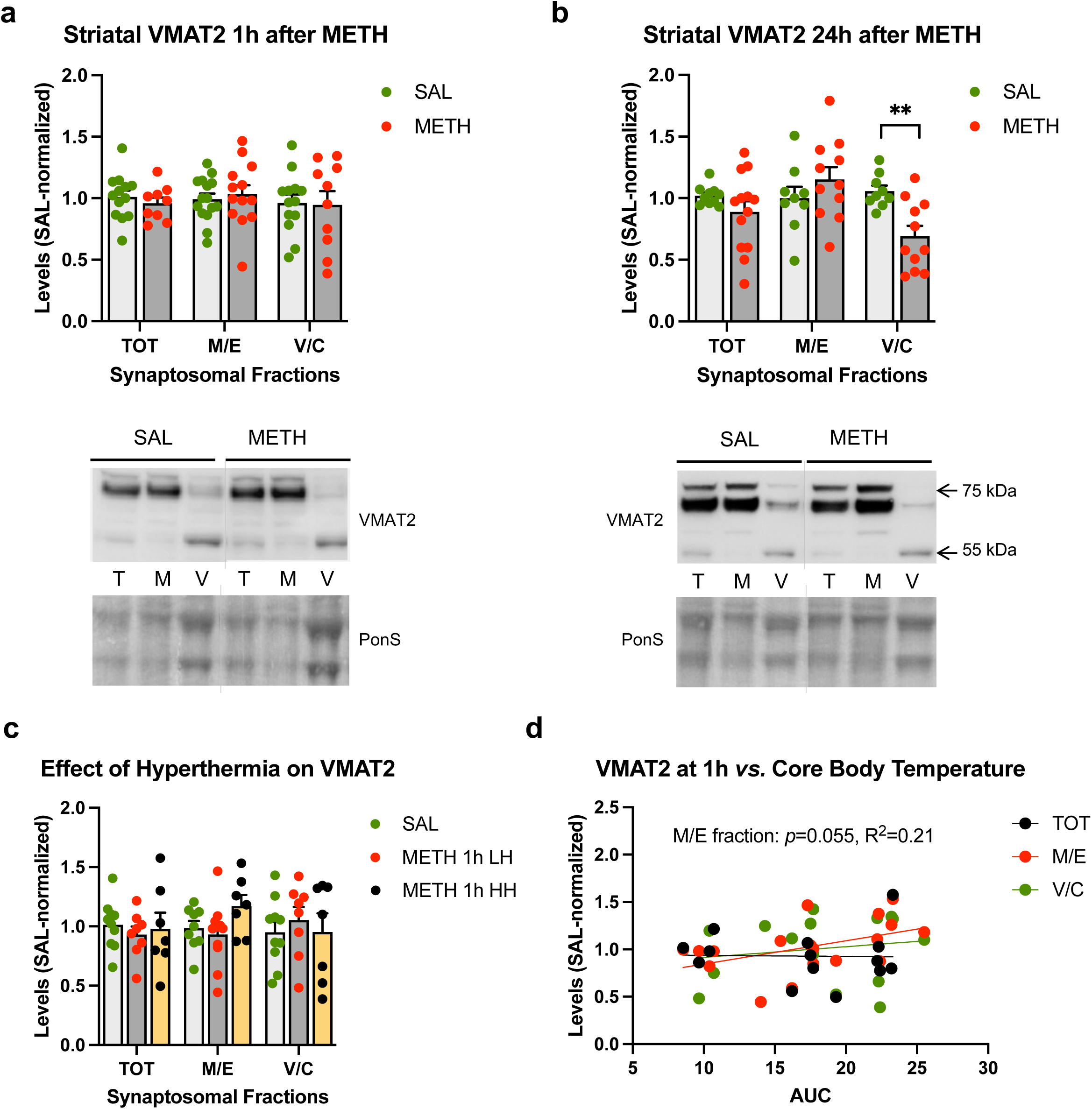
The effects of 4x8mg/kg METH binge on VMAT2 immunoreactivity in striatal synaptosomal fractions. Immunoreactivity of VMAT2 in total (TOT or T), membrane/endosomal (M/E or M), and vesicular/cytosolic (V/C or C) synaptosomal fraction of the striatum in rats euthanized at 1h **(a)** or 24h **(b)**. \*\**p*<0.01, n=9-11. **(c)** Immunoreactivity of VMAT2 in striatal synaptosomal fractions in high hyperthermia (HH) and low hyperthermia (LH) rats. **(d)** Correlations of VMAT2 immunoreactivity in synaptosomal fractions with core body temperature (area under the curve AUC) of rats sacrificed 1h after the last METH dose. A strong trend toward statistical significance was detected in membrane/endosomal fraction (*p*=0.055). Values are expressed as mean ± SEM. Vertical grey lines show where the blot was cut for rearrangement. Abbreviations: METH, methamphetamine; SAL, saline; PonS, Ponceau S.

### Parkin does not affect VMAT2 immunoreactivity in striatal synaptosomal fractions

Since parkin is associated with MTs and actin filaments ^46,47^ and the available evidence indicates its involvement in exocytosis processes, parkin may play a role in the intracellular trafficking of VMAT2-containing vesicles. To test this hypothesis, we assessed VMAT2 immunoreactivity in striatal synaptosomes from rats overexpressing parkin in the nigrostriatal DA pathway. Parkin overexpressed about 5-fold in all synaptosomal fractions (Fig.S2b and S3). Binge METH-treated parkin overexpressing rats displayed a similar profile of thermal response to METH as WT rats did (treatment *F*(1,10) = 102, *p*<0.0001, time (*F*(1.77,17.7) = 18.1, *p*<0.01, and treatment *x* time interaction (*F*(4,40) = 4.92, *p*<0.01, one-way RT ANOVA followed by Sidak *post hoc* test) (Fig.5a). As in the first METH experiment, binge METH treatment caused a small decrease in parkin immunoreactivity in WT rat striatum that displayed a trend toward statistical significance when analyzed by Student’s *t*-test (-26%, *p*=0.09, n=5) (Fig.4b, a representative blot is presented in Suppl. Fig.S3). The trend was lost when WT and PO data were analyzed by two-way ANOVA. METH treatment did not significantly change parkin immunoreactivity in parkin overexpressing rats (treatment and treatment x parkin levels *p*>0.05; parkin levels *F*(1,16) = 31.4, *p*<0.0001) (Fig.5b). Overexpression of parkin did not affect VMAT2 immunoreactivity in any synaptosomal fraction in saline-treated rats (*p*>0.05, n=5, multiple unpaired t-tests with Holm-Sidak correction for multiple comparisons) (Fig.5c). Binge METH did not decrease VMAT2 immunoreactivity in striatal synaptosomal fractions from WT or parkin overexpressing rats at 1h after the last METH injection (*p*>0.02, n=5, wo-way ANOVA with Holm-Sidak *post hoc* test) (Fig.5d). There was no significant correlation between total parkin immunoreactivity and total, membrane/endosomal, or vesicular/cytosolic immunoreactivity of VMAT2 in METH-treated rats at 1h after the treatment (*p*>0.05, Pearson correlation test) (Fig.4e).

**Figure 5.**
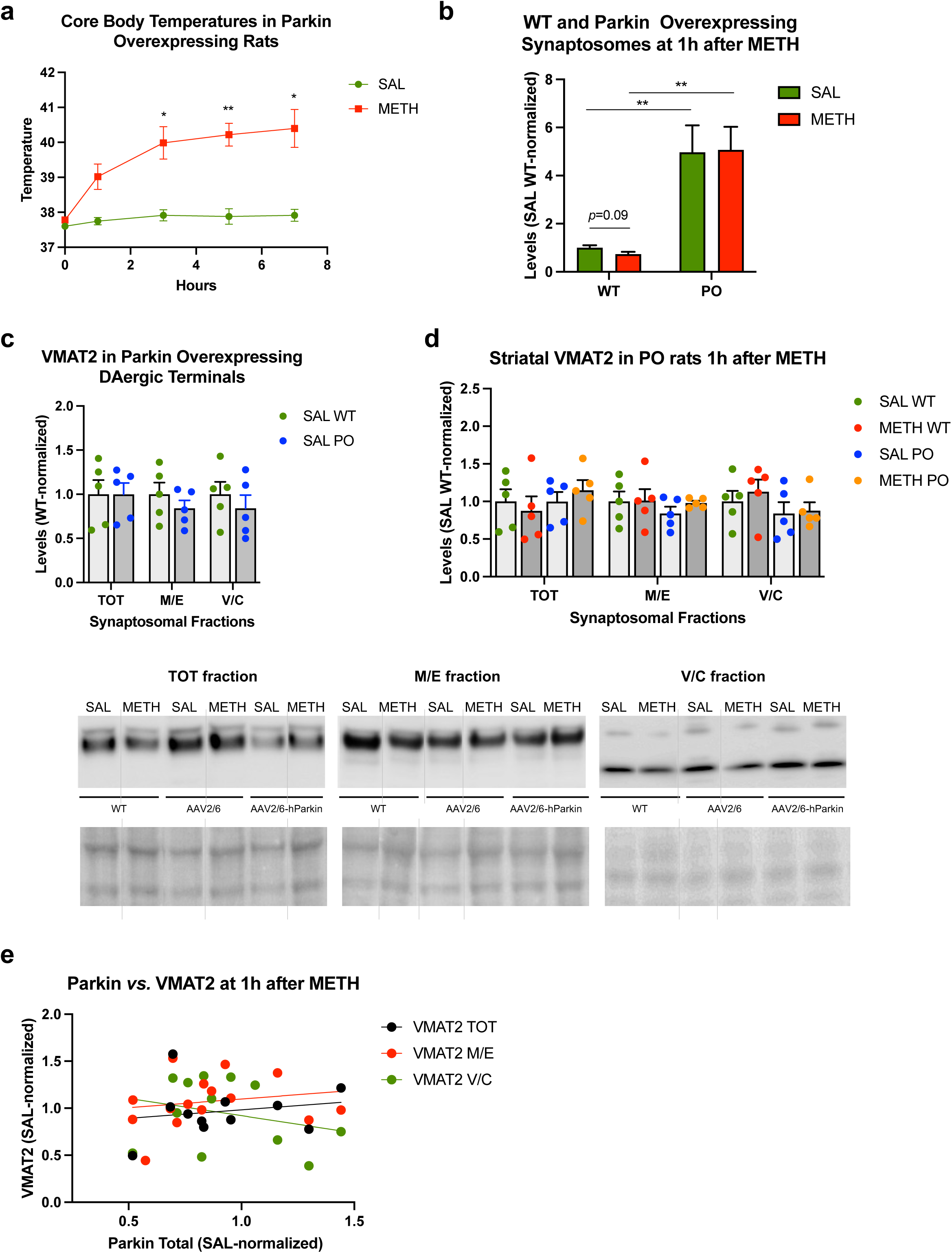
Parkin does not affect VMAT2 immunoreactivity in striatal synaptosomal fractions. **(a)** Core body temperatures (°C) of wild-type rats and rats overexpressing parkin in the nigrostriatal dopamine pathway. \**p*<0.05, \*\**p*<0.01. **(b)** Immunoreactivity of parkin in total synaptosomal fraction of the striatum in rats euthanized at 1h after METH or saline. \**p*<0.05, n=5. Parkin overexpression was about 5-fold in both treatment groups. METH did not significantly alter parkin immunoreactivity in the wild-type (*p*=0.09) or parkin overexpressing (*p*>0.1) striatal synaptosomes at 1h after the treatment. **(c)** Immunoreactivity of VMAT2 in total (TOT), membrane/endosomal (M/E), and vesicular/cytosolic (V/C) synaptosomal fraction in the striatum from wild-type and parkin overexpressing saline-treated rats. **(d)** Immunoreactivity of VMAT2 in striatal synaptosomal fractions in wild-type and parkin overexpressing rats treated with METH or saline at 1h after the treatment. **(e)** Correlations of VMAT2 immunoreactivity in synaptosomal fractions with total synaptosomal parkin immunoreactivity. Values are expressed as mean ± SEM. Vertical grey lines show where the blot was cut for rearrangement. Abbreviations: AAV2/6, adeno-associated viral vector 2/6; METH, methamphetamine; SAL, saline; PO, parkin overexpressing; WT, wild type; PonS, Ponceau S; AAV2/6-parkin, parkin-encoding AAV2/6; AAV6, non-coding AAV2/6.

### The effects of 4x8mg/kg METH binge on CDCrel-1 immunoreactivity in striatal synaptosomal fractions

To determine the effect of METH binge on CDCrel-1 levels in rat striatal synaptosomes, we assessed CDCrel-1 immunoreactivity in the WT rats treated with saline or 4x8mg/kg binge METH. As compared to saline controls, no significant changes in CDCrel-1 immunoreactivity in striatal synaptosomal fractions were detected in WT rats at 1h after the last METH injection (Fig.6a) whereas a significant decrease in CDCrel-1 immunoreactivity was detected in total and vesicular/cytosolic fraction at 24h after METH as compared to saline controls (-13% and -22%, respectively, *p*<0.05, n=9-10, multiple unpaired *t*-tests with Holm-Sidak correction for multiple comparisons) (Fig.6b). As with other measured indices, a wide range of individual responses to the drug was observed at 1h, with some rats having very high CDCrel-1 immunoreactivity (Fig.6a). After dividing the METH group into two subgroups (group 1 with total synaptosomal CDCrel-1 immunoreactivities at >120% and group 2 with total synaptosomal CDCrel-1 immunoreactivity at <120% increase, no change or a decrease), significant METH-induced increases in total and membrane/endosomal fractions in group 1 were observed (+58%, *p*<0.0001 and +55%, *p*<0.001, n=6-13, multiple unpaired *t*-tests with Holm-Sidak correction for multiple comparisons) and significant METH-induced decreases in total and vesicular/cytosolic fractions in group 2 emerged (-10% and -21%, respectively, *p*<0.05, n=11-13, multiple unpaired *t*-tests with Holm-Sidak correction for multiple comparisons) (Fig.6c, d). CDCrel-1 immunoreactivity was not significantly changed or markedly varied in the control cerebellum (Suppl. Fig.S1). After dividing the METH rats according to their thermal responses to METH (low or high hyperthermia: LH or HH), a small statistically significant decrease in CDCrel-1 immunoreactivity was observed in the vesicular/cytosolic fraction in the HH rats as compared to the controls (-22%, *p*<0.05, multiple unpaired *t*-tests with Holm-Sidak correction for multiple comparisons) (Fig.6e). There was no significant correlation between the core body temperature and CDCrel-1 immunoreactivity in synaptosomal fractions, suggesting that core body temperature does not have marked influence on CDCrel-1 levels in striatal synaptosomes (*p*>0.05, Pearson correlation test) (Fig.4f).

**Figure 6.**
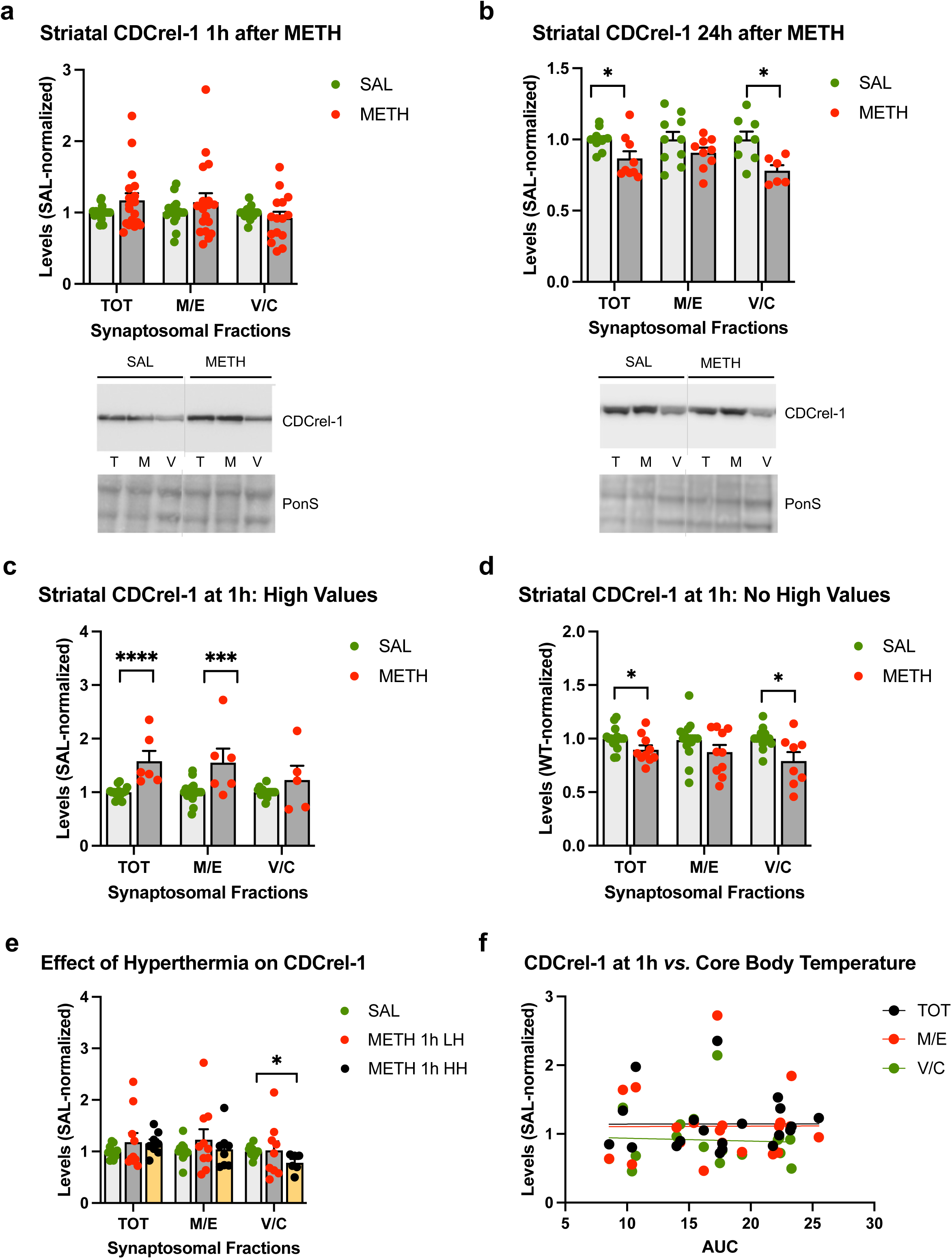
The effects of 4x8mg/kg METH binge on CDCrel-1 immunoreactivity in striatal synaptosomal fractions. Immunoreactivity of CDCrel-1 in total (TOT or T), membrane/endosomal (M/E or M), and vesicular/cytosolic (V/C or C) synaptosomal fraction of the striatum in rats euthanized at 1h **(a)** or 24h **(b)**. \**p*<0.05, n=9-10. **(c)** Immunoreactivity of CDCrel-1 in striatal synaptosomal fractions separated into two subgroups based on individual variability to METH. \**p*<0.05, \*\*\**p*<0.001, \*\*\*\**p*<0.0001, n=6-13. **(e)** Immunoreactivity of CDcrel-1 in striatal synaptosomal fractions in high hyperthermia (HH) and low hyperthermia (LH) rats. \**p*<0.05, n=6-10. **(f)** Correlations of CDCrel-1 immunoreactivity in synaptosomal fractions with core body temperature (area under the curve AUC) of wild-type rats sacrificed 1h after the last METH dose. Values are expressed as mean ± SEM. Vertical grey lines show where the blot was cut for rearrangement. Abbreviations: METH, methamphetamine; SAL, saline; PonS, Ponceau S.

### CDCrel-1 interactions with parkin in striatal synaptosomal fractions

*In vitro*, parkin regulates the levels of CDCrel-1, a protein found to regulate cytoskeleton organization and inhibit exocytosis ^28,32^. CDCrel-1 also induces neurodegeneration of DA neurons when overexpressed in the nigrostriatal DA pathway ^34^. Consequently, the METH-induced deficit in parkin could have resulted in increased CDCrel-1 levels and contributed to METH-induced neurodegeneration of DAergic terminals in the rat striatum at least in some of the rats. To determine the effect of parkin on CDCrel-1 levels in rat striatal synaptosomes, we first performed a co-immunoprecipitation experiment. As shown in Figure 6a, the anti-parkin antibody co-immunoprecipitated a small amount of CDCrel-1 with parkin in striatal synaptosomes from WT rats. Next, we assessed CDCrel-1 immunoreactivity in parkin overexpressing rats. Overexpression of parkin in the nigrostriatal DA pathway resulted in trends toward statistical significance in total and vesicular/cytosolic fractions in METH-naïve rats (-18%, *p*=0.09 and -43%, *p*=0.09, n=5, multiple unpaired *t*-tests with Holm-Sidak correction for multiple comparisons) (Fig.7b). CDCrel-1 immunoreactivity showed a trend toward statistically significant decrease in vesicular/cytosolic fraction after binge METH administration in WT rats (-33%, *p*=0.079, n=5, multiple unpaired *t*-tests with Holm-Sidak correction for multiple comparisons) (Fig.7b). The trend towards statistical significance was lost when all treatment data was analyzed by two-way ANOVA (Fig.7c). No METH-induced high CDCrel-1 immunoreactivities were observed in this experiment, likely because of the smaller sample size as compared to the first experiment with WT rats. These results agree with those shown in Figure 7d (a subgroup with no high CDCrel-1 values). As aforementioned, METH-naïve parkin overexpressing rats displayed a 33% decrease in vesicular/cytosolic CDCrel-1 as compared to WT rats. Administration of METH binge did not further decrease CDCrel-1 immunoreactivity in this fraction in PO rats (Fig.7c). Correlation of parkin immunoreactivity with CDCrel-1 immunoreactivity was positive ad statistically significant in total synaptosomal fraction at 1h after METH (*p*<0.05, R^2^=0.259, Pearson correlation test) (Fig.7d).

**Figure 7.**
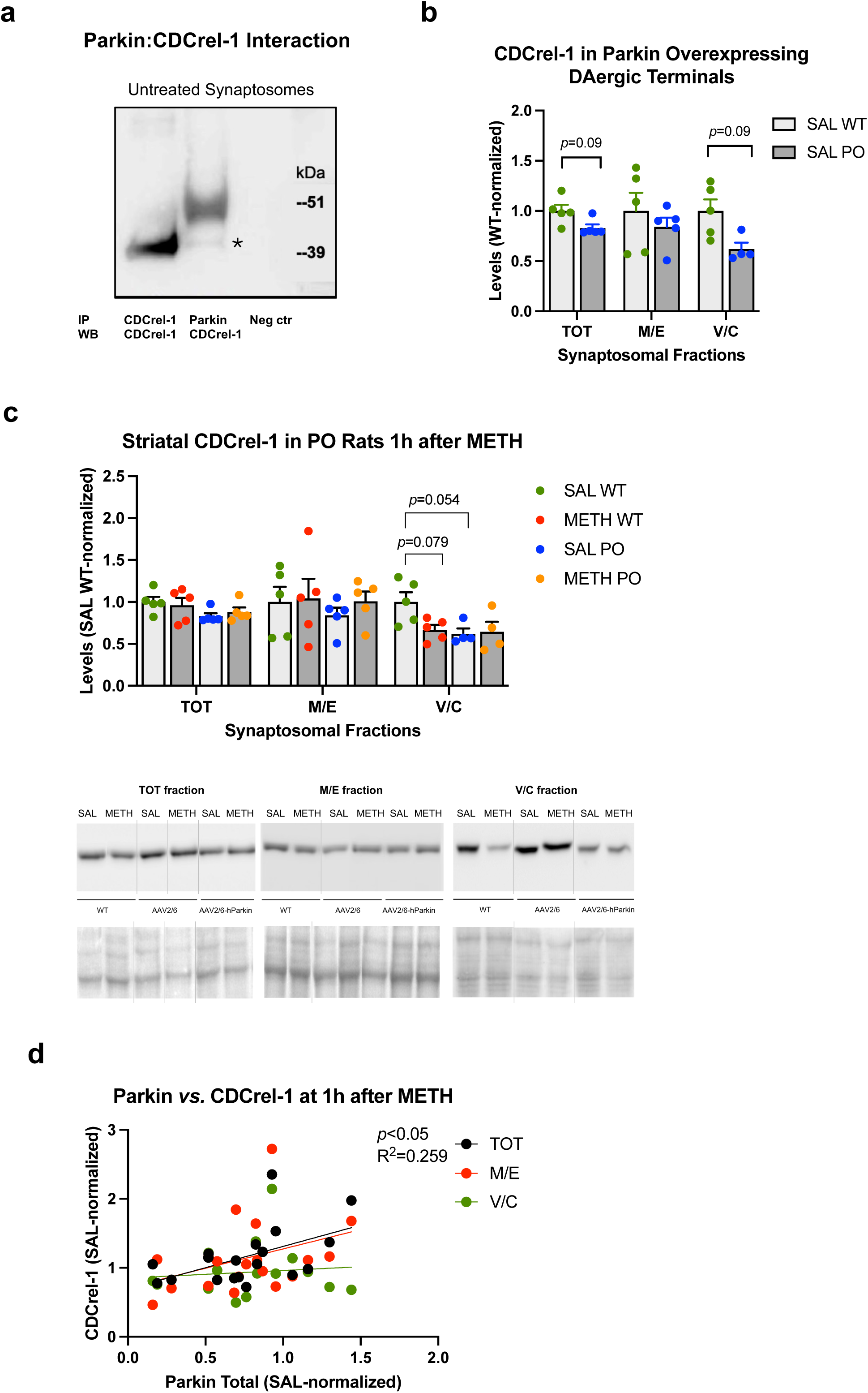
CDCrel-1 interactions with parkin in striatal synaptosomal fractions. **(a)** Anti-parkin antibody immunoprecipitated CDCrel-1 from untreated striatal synaptosomes (asterisk). **(b)** There was a weak trend toward statistical significance for the overexpression of parkin in the nigrostriatal dopamine pathway decreasing CDCrel-1 levels in total and vesicular/cytosolic synaptosomal fraction (*p*=0.09, n=5). **(c)** Trends toward statistical significance for parkin and METH decreasing CDCrel-1 immunoreactivity in wild-type rats at 1h after the last dose of the drug were detected (*p*=0.054 and *p*=0.079, respectively, n=5). METH treatment did not decrease CDcrel-1 levels in parkin overexpressing rats. **(d)** Correlations of CDCrel-1 immunoreactivity in synaptosomal fractions with parkin immunoreactivity of wild-type rats sacrificed 1h after the last METH dose. A statistically significant positive correlation was found in the total synaptosomal fraction. Values are expressed as mean ± SEM. Abbreviations: METH, methamphetamine; SAL, saline; PonS, Ponceau S; AAV2/6-parkin, parkin-encoding AAV2/6; AAV6, non-coding AAV2/6; IP, immunoprecipitation; WB, western blotting.

### CDCrel-1 interactions with VMAT2 in striatal synaptosomes

We next examined whether there is CDCrel-1: VMAT2 interaction in striatal synaptosomes in METH-naïve and METH-exposed rats. Anti-VMAT2 antibody immunoprecipitated CDCrel-1 from untreated synaptosomes and METH-treated synaptosomes, with more CDCrel-1 being pulled down at 1h after METH treatment (Fig.8a, b). Furthermore, there was a significant correlation between CDCrel-1 and VMAT2 immunoreactivity in membrane/endosomal fraction (Fig.8c).

**Figure 8.**
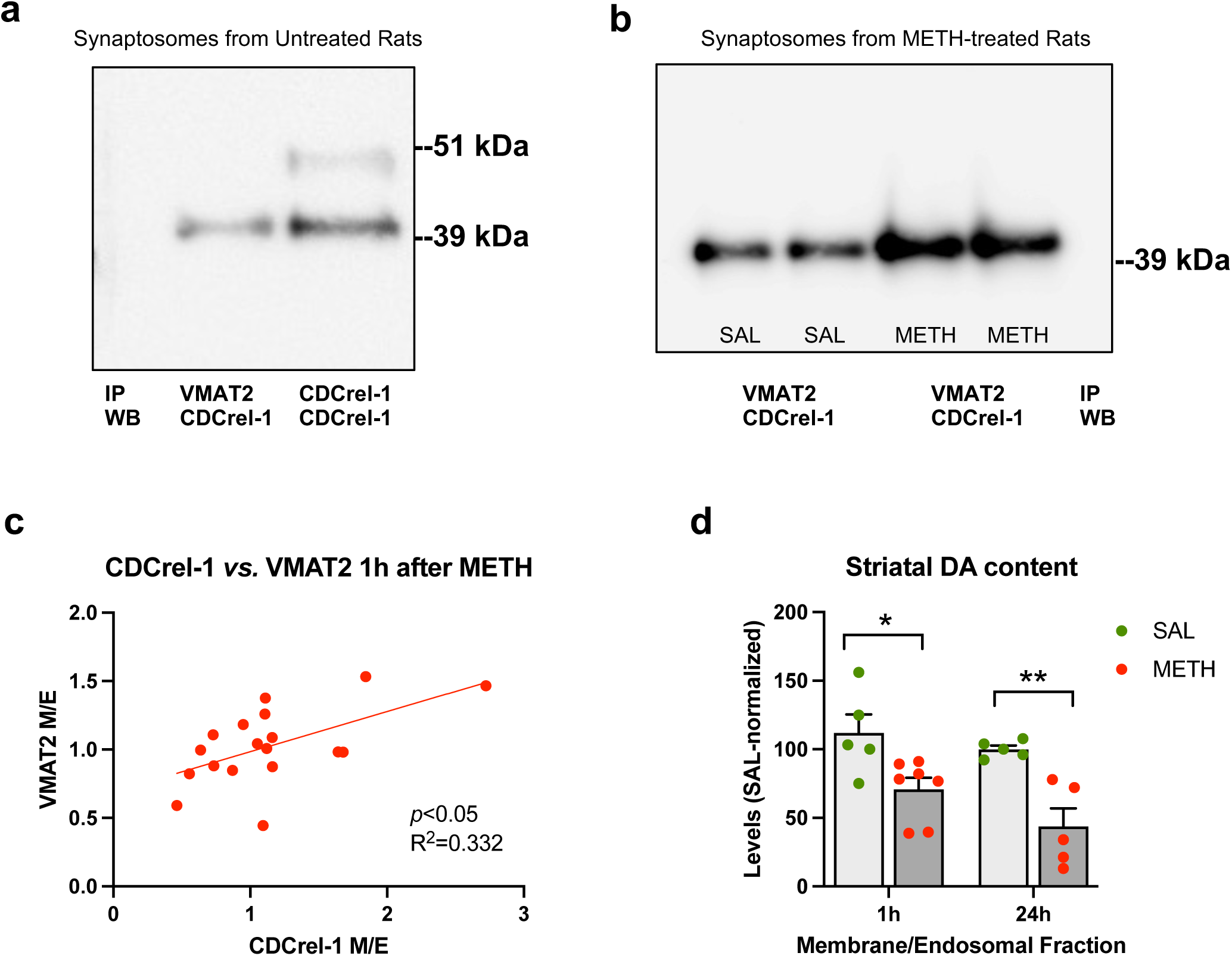
CDCrel-1 interactions with VMAT2 in striatal synaptosomes. **(a)** Anti-VMAT2 antibody immunoprecipitated CDCrel-1 from untreated striatal synaptosomes. **(b)** METH treatment increased the amount of immunoprecipitated CDCrel-1 at 1h after the last dose of the drug. **(c)** There was a significant positive correlation between CDCrel-1 and VMAT2 immunoreactivity in membrane/endosomal fraction of striatal synaptosomes (*p*<0.05). **(d)** Striatal VMAT2 vesicles associated with membrane/endosomal fraction had significantly lower dopamine content in METH-treated rats than saline-treated rats at 1h after the treatment (-29%). Over the next 24h dopamine content decreased to 66%. \**p*<0.05, \*\**p*<0.01, n=5-7. Values are expressed as mean ± SEM. Abbreviations: DA, dopamine; METH, methamphetamine; SAL, saline; IP, immunoprecipitation; WB, western blotting.

### The effects of 4x8mg/kg METH binge on dopamine levels in synaptosomal membrane/endosomal fraction

Neurotoxic binge METH largely depletes DA from all VMAT2 vesicle pools in striatal synaptosomes by 1h. To assess how much DA remained in membrane-bound VMAt2 vesicles, we assessed DA content in synaptosomal membrane/endosomal fractions at 1h after the last dose of METH or saline. There was a significant main effect of treatment (METH *vs.* saline) but no significant main effect of time (1h *vs.* 24h), on DA content in membrane/endosomal fraction (*F*(1,18) = 22.3, *p*<0.001, n=5-7, two-way ANOVA with Holm-Sidak *post hoc* test). DA content was decreased by 29% and 66% at 1h and 24h post-METH, respectively (Fig.8d).

### The effects of 4x8mg/kg METH binge on VMAT2-associated proteins in synaptosomal membrane/endosomal fraction

Intracellular transport of vesicles within terminals is mediated not only by MTs but also by actin microfilaments ^48^. Both MTs and actin filaments, as well as filamentous CDCrel-1, play essential roles in the neuronal exocytosis/endocytosis cycle. To examine whether METH binge alters exocytic and endocytic proteins associated with VMAT2 vesicles, we immunoprecipitated VMAT2 and its interacting partners from membrane/endosomal synaptosomal fractions of saline- and METH-treated rats sacrificed at 1h after their treatment and analyzed the immunoprecipitated proteins by mass spectrometry (n=4). Among 1281 proteins detected, the abundance of 10 proteins was significantly altered at FRD<0.15: 9 increased and 1 decreased (Table 1). Six of the 10 proteins were proteins involved in axonal/intracellular transport or exocytosis/endocytosis: α-tubulin N-acetyltransferase 1 (ATAT1), protein kinase C and casein kinase substrate in neurons protein 1 (PACN1, syndapin-1), MT-associated protein RP/EB family member 2 (MARE2), receptor-type tyrosine-protein phosphatase-like N (PTPRN), glia maturation factor beta (GMFB), and sorting nexin-17 (SNX17). ATAT localizes to clathrin-coated pits and acetylates α-tubulin on MTs at lysine 40. ATAT1 promotes MT destabilization and accelerates MT dynamics; this activity may be independent of acetylation activity ^49^. Syndapin-1 plays a role in the reorganization of the MT cytoskeleton via its interaction with MT-associated protein tau (MAPT) ^50^. This interaction decreases MT stability and inhibits MAPT-induced MT polymerization. Syndapin-1 also plays a role in the reorganization of the actin cytoskeleton and, therefore, is required for normal intracellular transport and synaptic vesicle endocytosis. The function of MARE2 is unknown. It may be involved in MT polymerization and dynamics ^51,52^. PTPRN is a transmembrane tyrosine phosphatase localized to dense-core synaptic vesicles ^53^. It may play a role in neurotransmitters loading into these vesicles. GMFB plays an important role in the growth, differentiation, and stress responses in neurons as well as glia ^54^. It may play a role in regeneration of axons and modulation of actin filaments ^55^. Nexin-17 is a member of the sorting nexin family localized to early endosomes ^56^ and regulates endocytic trafficking of several proteins ^57^. Other significantly changed proteins included 2 mitochondrial proteins: pyruvate dehydrogenase (acetyl-transferring) kinase isozyme 1 (PDK1) and mitochondrial coenzyme A transporter (SLC25A42), and 2 proteins involved in responses to stress: apolipoprotein D (APOD), and fibrinogen α-chain (FIBA). PDK1 is a kinase that plays a key role in the regulation of glucose and fatty acid metabolism and responses to cellular stresses ^58^. SLC25A42 transports coenzyme A into mitochondria in exchange for ADP. Both proteins could have co-immunoprecipitated with VMAT2 via association with the cytoskeleton and plasma membrane. APOD is a member of the lipocalin superfamily involved in lipid trafficking, food intake, inflammation, and antioxidative responses ^59^. FIBA is a blood-borne glycoprotein that has been found in neurons and glia ^60^. It functions during the early stages of wound repair. Ten differentially expressed proteins are presented in Table 1.

**Table 1.** List of differentially changed (FDR<0.15) proteins by binge METH treatment.

| Accession # | Protein Symbol | Protein Name | Function | Fold Change |
| --- | --- | --- | --- | --- |
| Q6MG11 | ATAT_RAT | Alpha-tubulin N-acetyltransferase 1 | Acetylates α-tubulin on MTs, promotes MT destabilization and accelerates MT dynamics | 9.2 |
| Q9Z0W5 | PACN1_RAT | Protein kinase C and casein kinase substrate in neurons protein 1 | Reorganization of MT and actin cytoskeleton, decreases MT polymerization and stability, required for bulk endocytosis | 2.7 |
| Q3B8Q0 | MARE2_RAT | Microtubule-associated protein RP/EB family member 2 | Unknown, may be involved in MT polymerization and dynamics | 1.9 |
| Q63259 | PTPRN_RAT | Receptor-type tyrosine-protein phosphatase-like N | Neurotransmitter loading into dense-core synaptic vesicles | 2.4 |
| Q63228 | GMFB_RAT | Glia maturation factor beta | Role in the growth, differentiation, and stress responses in neurons and glia, and actin-mediated endocytosis | 11 |
| Q6AYS6 | SNX17_RAT | Sorting nexin-17 | Regulates endocytic trafficking of several proteins, role in protein sorting and autophagy | 0.45 |
| Q63065 | PDK1_RAT | Pyruvate dehydrogenase (acetyl-transferring) kinase isozyme 1, mitochondrial | Regulation of glucose and fatty acid metabolism, and responses to cellular stresses | 3.3 |
| P0C546 | S2542_RAT | Mitochondrial coenzyme A transporter SLC25A42 | Transports coenzyme A into mitochondria in exchange for ADP | 2.0 |
| P06399 | FIBA_RAT | Fibrinogen alpha chain | Role in responses to injury and inflammation, and axonal repair | 2.5 |
| P23593 | APOD_RAT | Apolipoprotein D | Role in lipid trafficking, food intake, inflammation, and antioxidative responses, axon regeneration | 5.8 |

## Discussion

The major findings of this study are: (1) CDCrel-1 interacts with VMAT2 and has a role in VMAT2 trafficking under METH-induces stress, (2) there are wide individual differences in outbred Sprague Dawley rats in responses to METH neurotoxic effects such as hyperthermia and oxidative stress, (3) parkin does not have a major role in the regulation of VMAT2 or CDCrel-1 levels or trafficking in striatal neuronal terminals, and (4) changes in exocytosis/endocytosis cycle proteins are early responses to METH binge.

The Sprague Dawley rats employed in our study were outbred but were of the same strain, sex, and age, and they were housed and assessed under the same conditions. Therefore, the wide individual differences in thermal and molecular responses to METH binge were likely due to genetic differences such as variations in genes encoding thermal receptors, antioxidant enzymes, VMAT2, or dopamine transporter (higher function or levels of dopamine transporters at the membrane, more METH gets into DAergic terminals). Most studies on METH neurotoxicity used 4x10mg/kg METH (free base). The results from these studies were, for the most part, consistent. Therefore, the individual differences emerge at 4x8mg/kg and lower METH doses, or they emerge when the sample sizes are sufficiently increased (our sample sizes of the 1h cohorts were larger than the sample sizes employed in previous studies on METH neurotoxicity). In support of the latter scenario, we found less variability in parkin and CDCrel-1 immunoreactivity in the second experiment which employed five rats per group (WT *vs.* PO rat experiment, saline *vs*. METH). The observed U-shape-like responses to METH suggest that some rats could still upregulate their antioxidant mechanisms while some could not after exposure to 4x8mg/kg METH binge.

We previously determined that 4x10mg/kg METH decreased parkin levels in striatal synaptosomes at 1h and 24h after the last dose of the drug, with parkin levels returning to the baseline by 48h ^38^. The decrease was caused by oxidative modification and degradation of parkin. In this study, 4x8mg/kg METH decreased parkin levels in total and vesicular/cytosolic synaptosomal fractions at 1h but not at 24h after METH, suggesting that oxidative stress was less severe in the present study, allowing parkin levels to recover within 24h. The recovery could have been due to *de novo* synthesis of parkin and axonal transport to the terminals ^61^. As with body temperature, high variability in striatal parkin levels was observed in METH-treated rats, a decrease in some rats and no change, or even an increase, in others. Interestingly, significant parkin deficit was detected in rats with low hyperthermia with highly hyperthermic rats displaying mostly unchanged parkin levels. Hyperthermia itself can cause protein aggregation and oxidative stress ^62^, which in turn can decrease the number of viable MTs and inhibit axonal transport ^63^.

Under conditions of mild oxidative stress oxidized proteins are formed, but normally they are rapidly degraded by the 20S proteasome ^64–66^. Parkin is sensitive to DA-mediated oxidative modifications, and prone to misfolding and aggregating under oxidative stress ^38,67–70^. The 26S proteasome is also sensitive to oxidative stress, more so than the 20S proteasome ^71^. The 26S proteasome plays a predominant role in the normal turnover of parkin whereas the 20S proteasome (or lysosome) is more important for degradation of oxidized and aggregated parkin ^38,67,72^. When oxidative stress becomes severe, proteasomal function decreases either because of direct oxidative damage, because the accumulation of oxidized proteins exceeds the capacity of proteasomes to clear them, or because proteins are so severely altered that they are no longer recognized as substrates ^65,71^. In our previous study, we found 26S proteasome to be decreased at 1h after 4x10mg/kg binge METH ^31^. Overall, the data suggest that oxidatively modified parkin was successfully removed from striatal DAergic terminals in rats with low hyperthermia but not in highly hyperthermic rats, in which severe oxidative stress impaired the proteasome, lysosome, and retrograde axonal transport, which is also involved in the removal of damaged proteins ^73^. Another possibility is that oxidative stress changed parkin conformation obscuring the antibody-binding epitope. Regardless of the mechanism, the decrease in parkin function likely resulted in the impairment of multiple neuronal processes within DAergic terminals.

It is well established that the levels of antioxidant defenses increase in response to mild oxidative stress and decrease in response to severe oxidative stress ^74–76^. Our finding of increased parkin levels in synaptosomes from METH neurotoxicity-resistant cerebellum agrees with this evidence and suggests that a similar compensatory increase in parkin levels could have taken place in non-DAergic synaptosomes, which are in majority in striatal synaptosomes, masking parkin deficits and adding to the confounding effect of hyperthermia.

There are 3 pools of VMAT2 vesicles in the DAergic presynaptic terminal: (1) the reserve pool which is away from the active zone at the plasma membrane, (2) the proximal recycling pool that is closer to the active zone, and (3) the easily releasable pool of VMAT2 vesicles that are docked and ready for exocytosis ^77^. The vesicles are transported into terminals via anterograde axonal transport and can be removed from the terminals via retrograde transport. It has been reported that METH toxicity is associated with impaired VMAT2 trafficking in striatal synaptosomes at 1h and VMAT2 deficit at 24h after METH binge ^17,18,36,37^. Specifically, decreases in VMAT2 were reported in the vesicular/cytosolic fraction at 1h after METH, with membrane VMAT2 levels reported to be increased by one study in mice and not changed by the study in rats ^17,18^. These results suggested degradation of VMAT2 or its mobilization to a non-synaptosomal compartment, potentially by retrograde transport of damaged VMAT2 to the cell bodies in the substantia nigra *pars compacta*. In this study, a significant decrease in vesicular/cytosolic VMAT2 was detected only at 24h. Given that we used a lower dose of METH binge in this study than the other studies used (4x8 mg/kg vs. 4x10 mg/kg), METH-induced DA-mediated oxidative stress likely needed more time to rise to the threshold necessary to damage VMAT2, and decreased VMAT2 levels in the cytoplasm later than 1h after METH binge. Since there was high variability in VMAT2 levels in METH rats, an alternative explanation for the lack of expected VMAT2 deficit at 1h is an adaptive increase in the VMAT2 levels in some rats and a decrease in others. Despite unchanged levels, VMAT2 function was likely impaired by nitrosylation at 1h time point. Yamamoto group demonstrated modification of VMAT2 by nitrosylation at 1h after 4x10 mg/kg METH binge ^36^. No significant changes in VMAT2 levels were detected in high-hyperthermia or low-hyperthermia rats but a positive correlation of membrane/endosomal VMAT2 with core body temperature was observed in rats with high hyperthermia. This suggests retention or mobilization of VMAT2 vesicles to the plasma membrane in high-hyperthermia rats.

Parkin could influence axonal VMAT2 vesicle trafficking by its known interaction with MTs and their stabilization ^46^. Moreover, parkin could influence VMAT2 vesicle trafficking between their pools via interaction with α-synuclein ^21,78^, actin ^47^, and CDCrel-1, which was reported to be a substrate for parkin in cultured cells ^28^. CDCrel-1 is a filamentous protein found attached to membrane and synaptic vesicles ^79^. CDCrel-1 binds to syntaxin-1, which is a part of the soluble NSF attachment protein (SNAP) Receptor complex (SNARE), a complex crucial in exocytosis. By this interaction, CDCrel-1 may prevent the docking of vesicles to the membrane and decrease exocytosis ^32^. We expected that METH neurotoxicity would decrease parkin levels with a consequent increase in CDCrel-1 levels and altered distribution of VMAT2 vesicles between the vesicular/cytosolic and membrane/endosomal fraction in striatal synaptosomes. We found evidence to the contrary, suggesting that parkin does not significantly affect VMAT2 levels or its intracellular trafficking *in vivo*. This conclusion is supported by our previous finding of unchanged VMAT2 levels in the METH-naïve striatum in parkin knockout rats ^80^.

CDCrel-1 was reported to be a substrate for parkin *in vitro* ^28^. We detected only a small effect of parkin on CDCrel-1 levels in striatal synaptosomes in METH-naïve rats. This finding suggests that parkin is not a major E3 ligase degrading CDCrel-1 in rat nigrostriatal pathway and agrees with the finding of unchanged CDCrel-1 levels in the striatum of parkin knockout mice ^81^. Alternatively, as our synaptosomal preparations contained non-DAergic synaptosomes which also express CDCrel-1, the effect of parkin on CDCrel-1 could have been “diluted” in DAergic synaptosomes. METH binge had a variable effect on striatal synaptosomal CDCrel-1 levels at 1h. Presently, it is unclear why some rats displayed significant increases in CDCrel-1 levels in striatal synaptosomes while others displayed significant decreases (U-shaped response to METH). Since most of the rats with high CDCrel-1 immunoreactivities fell into the LH group (Fig.6e), the reason might be low levels of parkin and decreased degradation of CDcrel-1 in this fraction in LH rats (Fig.3c). This variability did not depend on hyperthermia or parkin levels although high CDCrel-1 levels appeared to be accompanied by high parkin levels in METH rats in membrane/endosomal fraction. Of note, CDCrel-1 levels correlated negatively with parkin levels in METH-treated rats when high CDCrel-1 data points were removed. METH binge caused a deficit in CDCrel-1 in WT rats at 1h, and parkin overexpression appeared to protect vesicular/cytosolic CDCrel-1 from being decreased by METH. These results suggest that parkin is not engaged in the degradation of CDCrel-1 shortly after METH insult and that METH damages both proteins by oxidative stress and alters their interaction. At 24h, CDCrel-1 levels were decreased in total and vesicular/cytosolic fraction, suggesting the removal of damaged CDCrel-1 from cytosol following METH. The most apparent interaction found in our study was that between CDCrel-1 and VMAT2. This interaction was higher in METH-treated rats as compared to saline-treated rats at 1h time point and significant in the membrane/endosomal fraction. This finding suggests that an increased number of CDCrel-1 filaments may have “entrapped” VMAT2 vesicle at the synaptosomal plasma membrane in some rats, inhibiting exocytosis. Such “entrapment” would also prevent VMAT2 vesicle recycling and proper sequestration of cytosolic DA. This notion agrees with previous findings. CDCrel-1 has been suggested to negatively regulate synaptic vesicle release at presynaptic terminals by forming filamentous barricades at the presynaptic membrane upon interaction with syntaxin-1, which parkin ubiquitinates ^32^. It is possible that, in some rats, parkin (or other E3 ligase) is severely damaged and aggregated in METH-exposed striatal terminals, not able to ubiquitinate CDCrel-1. This, together with decreased 26S activity ^31^, leads to CDCrel-1 accumulation, disabling the release of syntaxin-1 to enhance VMAT2 vesicles docking ^28^. In summary, multiple toxic effects of METH (hyperthermia, oxidative stress, inflammatory mediators, excitotoxicity) influence levels of parkin, VMAT2, and CDCrel-1 in striatal terminals at different degrees in different rats. This individual variability in responses to binge METH indicates variability in stress responses in Long Evans rats.

There is evidence for METH altering the exocytosis/endocytosis cycle. For example, functional studies showed that amphetamines increase endocytosis of dopamine transporter ^82^ and glutamatergic glutamate transporter EAAT3 ^83^ in DA neurons. Proteomic/genetic studies reported altered brain levels of proteins with functions related to the cytoskeleton, transport, endocytosis, and exocytosis after exposure to METH ^84,85^. Finally, oxidative stress was reported to decrease endocytosis in neurons depending on the stressor (reviewed in ^86^). We have demonstrated that binge METH administration decreases the function of 26S proteasome and the levels of parkin (^31^ and Fig.3a) at 1h after the last METH injection. The function of 26S proteasome is important for exocytosis/endocytosis and DAergic neurotransmission ^87^ and several of parkin substrates are involved in the mediation of exocytosis, namely α-synuclein ^21^, α- and β-tubulin ^22^, synaptotagmins IV and 11 ^23,24^, synphilin-1 ^25^, endophilin, dynamin, synaptojanin 1 ^26^, and CDCrel-1 ^27,28^. Our understanding of alterations in the exocytosis/endocytosis cycle in METH neurotoxicity is incomplete. To gain more insight into these alterations, we immunoprecipitated VMAT2 and associated proteins from striatal membrane/endosomal fractions of saline- and METH-treated rats sacrificed at 1h. Endocytosis/exocytosis-related parkin substrates mentioned above were found in the immunoprecipitates but not significantly changed by METH binge, likely due to small sample sizes (n=4). Nevertheless, ten proteins were differentially expressed by binge METH treatment at 1h: nine increased and one decreased in abundance. Six of these proteins are involved in axonal/intracellular transport and exocytosis/endocytosis: ATAT1, PACN1, (syndapin-1), MARE2, PTPRN, GMFB (increased), and SNX17 (decreased).

ATAT1 localizes to clathrin-coated pits and acetylates α-tubulin. It promotes MT destabilization and accelerates MT dynamics ^49^. Tubulin hyperacetylation is associated with cellular responses to stresses, including oxidative stress ^88^, and increased cell survival through the induction of autophagy ^89^. Protein syndapin-1 also decreases MT stability and is required for activity-dependent bulk endocytosis (ADBE) of synaptic vesicles ^50^. ADBE generates many synaptic vesicles during intense neuronal activity or increased temperature, replenishing the reserve pool of the vesicles ^90,91^. An increase in syndapin-1 levels suggests a need for new vesicles for the storage of DA released into the cytosol. PTPRN is an important transmembrane protein in dense-core synaptic vesicles (DCVs), involved in cargo loading and exocytosis of DA in PC12 cells ^53^. The function of MARE2 is unknown. Available evidence suggests it may play a role in MT dynamics ^51^ and axonal delivery of DCVs ^52^. Upregulation PTPRN and MARE2 suggests mobilization of DCVs to the plasma membrane of DAergic terminals. VMAT2 is associated mainly with small synaptic vesicles but it is also found in DCVs ^92^. Furthermore, DA was reported to be co-stored with cholecystokinin in DCVs ^93^. It can be envisioned that DCVs vesicles were mobilized from substantia nigra *pars compacta* cell bodies to striatal terminals to increase DA storage of and/or replace vesicles with dysfunctional nitrosylation-damaged VMAT2. GMFB is upregulated in several neuroinflammatory and neurodegenerative conditions ^54^ and is associated with actin-mediated endocytosis ^55^. Nexin-17 interacts with a multitude of receptors, including integrins, in a sequence-specific manner to regulate their recycling ^56,57,94^. Furthermore, nexin-17 participates in the endocytic trafficking and processing of potentially harmful proteins and is linked to autophagy ^56^. FIBA (fibrinogen α-chain) is a blood plasma protein reported to be expressed by astrocytes and neurons under neuroinflammatory or neurodegenerative conditions^60^. FIBA also interacts with its receptors on neurons as well as with integrins and may be involved in axonal repair ^60^. Both nexin-17 and FIBA bind to integrin α5β1 which is present on endothelial cells ^94,95^. This integrin is involved in inflammatory responses and neuronal regeneration ^96^. The proteomic results suggest that both proteins were connected to striatal axonal terminal membranes and involved in responses to METH-induced injury to these terminals and to the blood-brain barrier (BBB). Another protein involved in responses to stress that was found to increase was APOD. APOD is a member of the lipocalin superfamily involved in lipid trafficking, inflammation, and antioxidative responses e.g., APOD is involved in the axon regeneration process as a lipid transporter ^59^.

PDK1 and SLC25A42 are mitochondrial proteins. As both mitochondria and synaptic vesicles are attached to actin and plasma membrane; therefore, they have been pulled down by α-VMAT2 antibody via this connection. PDK1 is a kinase that inhibits the formation of acetyl-coenzyme A and decreases mitochondrial respiration. By this mechanism, it protects cells against oxidative stress and apoptosis ^58^. SLC25A42 transports coenzyme A into mitochondria in exchange for ADP. Coenzyme A is important for the Krebs cycle function and, consequently, for mitochondria function. Several steps in the exocytosis/endocytosis cycle require ATP, including ADBE which relies on ATP-dependent actin polymerization ^58^. Increases in both PDK1 and SLC25A42 would have opposing effects on mitochondrial function with a net result being decreased levels of ATP as it was found in deficit at 1h after binge METH ^97^. PDK1 levels likely increased in response to METH-induced oxidative stress and inflammation while SLC25A42 levels increased to provide ATP for ADBE.

Tubulins and actin are cytoskeletal proteins that are present in abundance in the cytosol and synaptic active zones ^98^ where they are involved in intra-neuronal transports and the exocytosis/endocytosis cycle. The deficits in β-actin immunoreactivity observed in all synaptosomal fractions at 1h after METH suggest a negative impact on intra-synaptosomal transports as well as exocytosis/endocytosis cycle of synaptic vesicles. The increase in GMFB suggests reduced actin polymerization at 1h after METH that occurred to combat METH toxicity. It has been observed that DAergic neurons and astrocytes in GMFβ^-/-^ mice have reduced sensitivity to oxidative stress ^55^. GMFB overexpression reduces the levels of actin-related protein 2/3 complex (Arp2/3), the facilitator of actin polymerization ^99^. METH enhances or decreased actin polymerization depending on the treatment regimen and can increase actin cycling ^43^. For example, METH treatment increased Arp2/3 and enhanced actin polymerization as well as endocytosis of protein occludin in endothelial cells. Arp2/3 inhibition reversed actin polymerization, increased occludin levels, and restored disrupted BBB ^43^. Our DA and VMAT2 data indicate that VMAT2 vesicles associated with the membrane/endosomal fraction in the METH-treated striatum have not changed in number but were partially depleted of DA at 1h.

The lack of changes in VMAT2 levels at 1h after METH does not exclude a scenario of an increased rate of VMAT2 between the plasma membrane and endosomes, or between the cell body and the terminal. Our proteomic data indirectly suggests an increased rate of endocytosis/exocytosis cycle of VMAT2 in METH-exposed striatal synaptosomes at 1h, which does not result in significant changes in VMAT levels in the studied fractions because the rates of removal equal the rates of replacement. The available evidence suggests that parkin is involved in VMAT2 synaptic vesicle cycling. Thus, Song and colleagues reported the impairment of the endocytic pathway with which nexin-17 is associated in parkin-deficient cells ^30^. A recent study by the same group demonstrated that parkin was recruited to synaptic vesicles in a neuronal activity-dependent manner to promote vesicle endocytosis in DA neurons whereas parkin loss of function resulted in accumulation of oxidized DA ^29^. Parkin deficit or excess could have influenced VMAT2 vesicle trafficking between the plasma membrane and endosomes which we would not detect because of the way we fractionated synaptosomes. The lack of influence of parkin overexpression on VMAT2 immunoreactivity in striatal synaptosomes suggests that parkin does not play a major role in parkin degradation or trafficking along MTs in striatal terminals.

In agreement with the previous studies, we found decreased immunoreactivity of VMAT2 in vesicular/cytosolic fraction at 24h after METH administration. This decrease was accompanied by decreased immunoreactivity of CDCrel-1, suggesting retrograde transport of damaged VMAT2 vesicles, and CDCrel-1, out of the terminals or the beginning of DAergic terminal neurodegeneration at this time point. In support of the first scenario, our group showed that anterograde transport is still dysfunctional at 3 days after binge METH administration; and therefore, not able to replenish DAergic terminals with newly synthesized proteins ^100^.

Overall, the results suggest that at 1h after METH binge, DAergic neurons are engaged in counteracting METH-induced toxic effects, including oxidative stress- and hyperthermia-induced inhibition of synaptic vesicle cycling, and include increased autophagy and endocytosis. Trafficking of parkin, VMAT2, and CDCrel-1 is altered differently between the rats, likely depending on the level of hyperthermia and oxidative stress each animal could combat based on genetic makeup. In conclusion, our results indicate that studying large groups of outbred rats such as can help define individual genetic and molecular differences in responses to METH neurotoxicity which, in turn, will aid treating humans suffering from METH use disorder and its neurological consequences.

## Acknowledgments

This work was supported by NIH grant DA023085. We thank Akhil Sharma and Bryan Killinger for their help with animal work. We acknowledge the assistance of the Wayne State University Proteomics Core that is supported through NIH grants P30ES036084, P30CA022453, and S10OD030484.

## Author Contributions

A.M. designed the experiments, analyzed the data, prepared the figures, and wrote the manuscript. H.C. conducted the experiments, quantified the data, and helped with manuscript preparation. P.S. and N.C. performed proteomic analysis. B.S. provided the AAV2/6 vectors. All authors reviewed the manuscript.

