## Supplementary Material for "Neurotoxic Methamphetamine Doses Alter CDCel-1 Levels and Its Interaction with Vesicular Monoamine Transporter-2 in Rat Striatum"

The file contains:

Figure S1

Figure S2

Figure S3

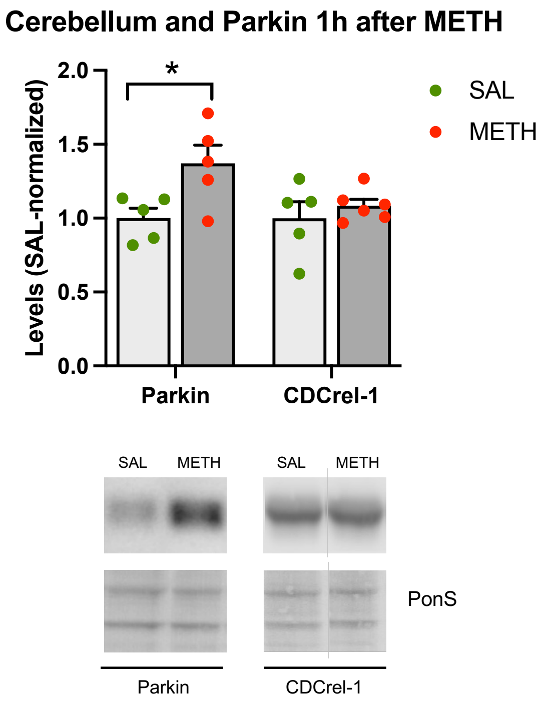


**Figure S1. The effects of 4x8mg/kg METH binge on parkin and CDCrel-1 immunoreactivity in cerebellar synaptosomes.** Immunoreactivity of parkin (left) and CDCrel-1 (right) in cerebellar synaptosomes in saline- and METH-treated rats at 1h after the treatment. *$p<0.05$. Vertical grey lines show where the blot was cut for rearrangement. Values are expressed as mean ± SEM. Abbreviations: METH, methamphetamine; SAL; PonS, Ponceau S.

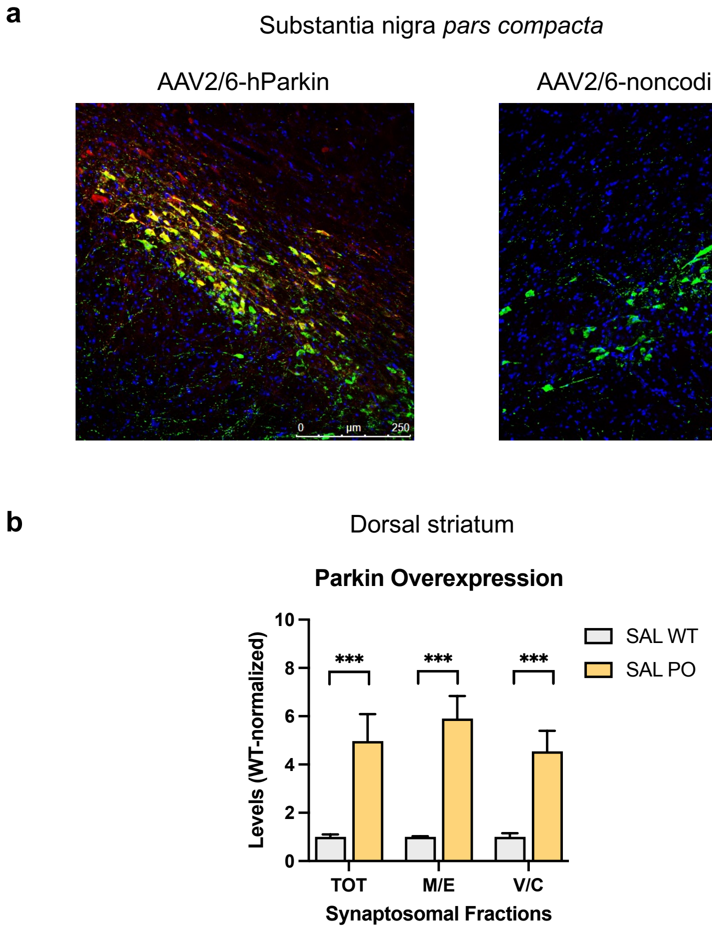


**Figure S2. Validation of parkin overexpression in the substantia nigra pars compacta and dorsal striatum. (a)** The left SNpc was microinjected with parkin-encoding AAV2/6 gene transfer vector (AAV2/6-parkin) whereas the right substantia nigra was microinjected with non-coding AAV2/6. Immunoreactivity of parkin (red fluorescence) and dopaminergic marker tyrosine hydroxylase (green fluorescence) in the left and right substantia nigra *pars compacta* (SNpc). The yellow color indicates parkin immunofluorescence overlapping with tyrosine hydroxylase fluorescence, thus confirming parkin overexpression in the left SNpc. Parkin is at very low levels in the nigrostriatal pathway; therefore, it is hard to detect by immunofluorescence in wild-type rats. Bars: 200µm. **(b)** Parkin immunoreactivity (chemiluminescence) increased ~5-fold in all synaptosomal fractions in the left dorsal striatum after microinjection of the AAV2/6-parkin into the left SNc. ***$p$<0.001. Abbreviations: METH, methamphetamine; SAL.

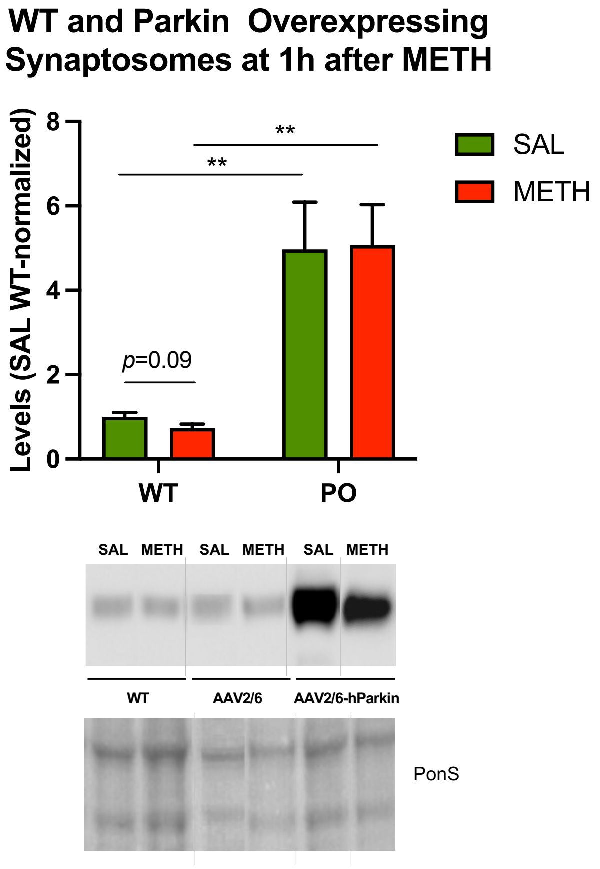


**Figure S3. The effects of 4x8mg/kg METH binge on parkin immunoreactivity in striatal synaptosomal fractions.** Figure 5b (top) shows parkin immunoreactivity in the total synaptosomal fraction in wild-type and parkin-overexpressing rats treated with saline or binge METH. At the bottom are representative blots and Ponceau loading control staining. ***p*<0.01. Vertical grey lines show where the blot was cut for rearrangement. Abbreviations: METH, methamphetamine; SAL.
